# AutoXAI4Omics: an Automated Explainable AI tool for Omics and tabular data

**DOI:** 10.1101/2024.03.25.586460

**Authors:** James Strudwick, Laura-Jayne Gardiner, Kate Denning-James, Niina Haiminen, Ashley Evans, Jennifer Kelly, Matthew Madgwick, Filippo Utro, Ed Seabolt, Christopher Gibson, Bharat Bedi, Daniel Clayton, Ciaron Howell, Laxmi Parida, Anna Paola Carrieri

**Author notes:** Correspondence should be addressed to Anna Paola Carrieri. Authors contributed equally to the manuscript (James Strudwick, Laura-Jayne Gardiner).

## Abstract

Machine learning (ML) methods have the potential of detailed insights of complex biological systems and today are increasingly used to analyse omics data for tasks such as the discovery of novel biomarkers and phenotype prediction. It can be extremely beneficial and powerful for scientists, domain experts, to easily run sophisticated, robust, and interpretable ML pipelines without the need for an in depth understanding of the code needed to train, tune, optimise ML algorithms. They can then focus on the biological interpretation and validation of the results and insights generated by ML models. Here, we present an entirely automated open-source explainable AI tool, AutoXAI4Omics, that performs classification and regression tasks from omics and tabular numerical data. AutoXAI4Omics accelerates scientific discovery by automating processes and decisions made by AI experts, e.g., selection of the best feature set, hyper-tuning of different ML algorithms and selection of the best ML model for a specific task and dataset. Prior to ML analysis AutoXAI4Omics incorporates feature filtering options that are tailored to specific omic data types. Moreover, the insights into the predictions that are provided by the tool through explainability analysis highlight associations between omic feature values and the targets under investigation e.g., predicted phenotypes, facilitating the discovery of actionable insights. AutoXAI4Omics is at: https://github.com/IBM/AutoXAI4Omics.

**Graphical Abstract:** 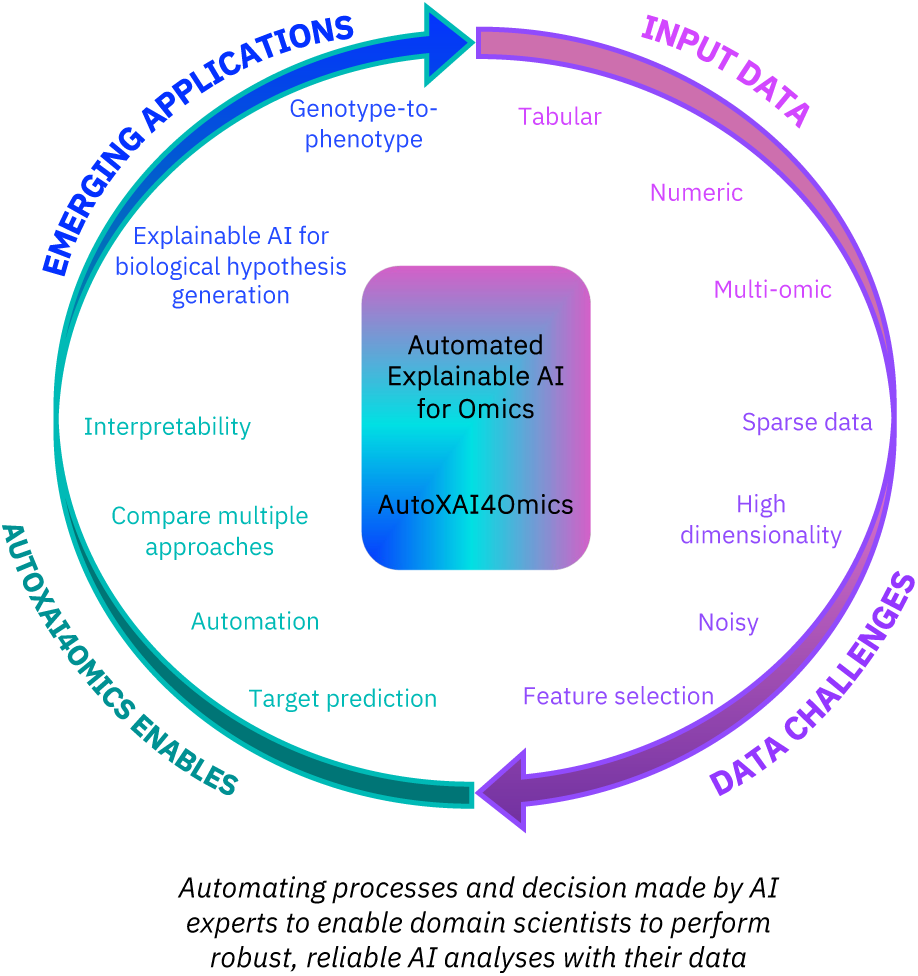

## Introduction

In recent years there has been an increase in utilising Artificial Intelligence (AI) and Machine Learning (ML) within scientific discovery and pipeline development for the analysis of healthcare and life science datasets [1]. AI represents the broader concept of simulating human intelligence by machines, while ML is a branch of AI commonly involving a computer learning autonomously. In supervised ML, it is the provision of labels for the input data that enables the learning; algorithms parse the input data (feature sets), learn patterns from it relating to the label (or target), and can then make predictions about the labels for unseen data. ML algorithms are well suited to large, heterogeneous, and complex input data sets such as omics (e.g. genomics, transcriptomics, proteomics, metabolomics, etc.) that are becoming increasingly common as the cost of sequencing decreases. In response to this, many available open-source AI and ML tools have been tailored to omics data [2] and related analytic questions that are commonly asked of such data sets. There is the need for research scientists to apply ML methods to perform prediction tasks from omics data and generate insights about the link between omics features and the target of investigation e.g., a phenotype, in a way that is quick, automated, and robust. However today, surprisingly, there are few entirely automated end-to-end open-source ML tools that can handle different types of omics data and perform generic classification or regression tasks.

### Current state of the art

Focusing on open-source offerings for omics data, there are a wide range of notable AI and ML tools that have been developed for specific use cases including; genotype-to-phenotype prediction (e.g., DeepCOMBI [3], AutoML-GWAS [4], Emedgene[5]), SNP calling (e.g., SNP-ML [6] and DeepVariant [7]), radiogenomics [8] (e.g., ImaGene [9]), omic feature selection (e.g., [10]), predicting gene regulation (e.g., BioAutoMATED [11], EUGENe [12], CRMnet [13]), single cell RNA gene regulatory analysis (e.g., scGeneRAI [14]), circadian rhythm detection (e.g., ZeitZeiger [15]) and protein folding prediction (e.g., Alphafold [16]). Many of these tools present a single ML method and focus on a specific data type or a few selected data types (e.g., genomics and transcriptomics). Furthermore, the majority of these tools are tailored to a single application domain e.g., for biomedical and drug discovery (summarised in the review by [2]) or plant science (summarised in the reviews by [17] and [18]).

It is not uncommon for both AI and ML models to be referred to as ‘black-boxes’, due to their complexity and lack of interpretability. This lack of interpretability (or explainability) can create concern for the user of the model, the domain expert, since it may allow issues and biases to be overlooked [19]. Consequently, Explainable AI (XAI) algorithms such as LIME [20] and SHAP [21] are applied to ML models to overcome these problems and improve trust in the models and their results by providing explanations of their predictions [22]. Furthermore, our previous work has harnessed such interpretability and explainability of models to gain biological insights into the target variable being predicted e.g., for prioritisation of key genes or markers associated with a predicted phenotype ([23], [24], [25]). This explainability goes beyond the interpretability offered by the feature importance metrics used in existing workflows since it directly explains the decisions made by a ML model [26, 27]. Finally, the problems of ’black-boxes’ can be accentuated by the lack of open-source pipelines, i.e., when a user is dependent on a model wrapped in a workflow that is also not transparent.

### Our offering: AutoXAI4Omics

In this manuscript we present our open-source, explainable, end-to-end ML tool for omics data (and any numerical tabular data generally). We call this software AutoXAI4Omics. AutoXAI4Omics, encompasses an automated workflow that takes the user from data ingestion through data pre-processing (filtering, normalisation, scaling) and automated feature selection (if desired) to generate a series of hyper-tuned optimal ML models for a given user-defined task (classification or regression). Figure 1 provides a high-level representation of the workflow. AutoXAI4Omics then aids the user in the evaluation of these models, suggesting the best model based on a range of key considerations (e.g., over-fitting and performance on held-out data) and generating a wide range of visualisations of the results. Finally, it facilitates interpretability of results, not only via feature importance inference, but using XAI to provide the user with a detailed global explanation of the most influential features contributing to each generated model. AutoXAI4Omics can process a large range of tabular data types including RNA-seq, microbiome-related data, metabolomics, genomics and other numerical tabular data e.g., clinical, environmental and demographic data. These data types can span domains such as human health and disease, drug discovery, plant phenotyping and environmental studies. It also has pre-processing capability tailored to omic data e.g., TMM normalisation for transcriptomics and common microbiome sample/feature filtering parameters.

**Figure 1:**
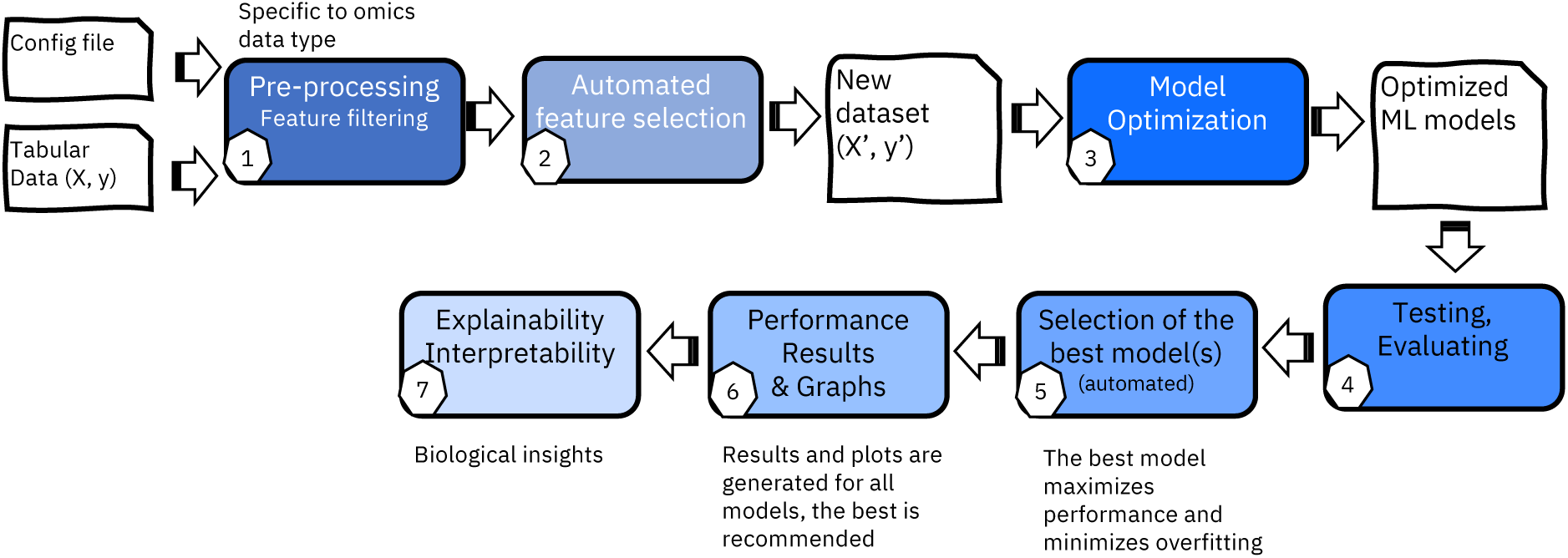
Overview of the AutoXAI4Omics XAI workflow from data input to results and interpretation. AutoXAI4Omics is available as a containerized command line tool where the intended audience is bioinformaticians or non-expert ML users generally. To use it the user provides an input data set (samples x features table), metadata (typically the target to predict by the ML) and a configuration file (config, json file). Our workflow advances existing open-source ML tools through our unique combination of having an end-to -end workflow (including omics-specific pre-processing, automated feature selection and best model recommendation), prioritising interpretability through XAI, being data type and application agnostic and being fully containerised to enable seamless installation and re-use. AutoXAI4Omics is available on Github at: https://github.com/IBM/AutoXAI4Omics.

In this manuscript we will go on to detail the options available to the user in AutoXAI4Omics and using three distinct biological case studies we will demonstrate and showcase its simplistic usage, performance, and application agnosticism. The various data sets and configuration files that we use here are also provided as supplementary material to encourage ease of re-use, since they can act as a template for new users. Our first biological use case focuses on the plant science domain where we predict if Barley accessions are two-rowed or six-rowed (relating to spikelet fertility), from genomic data, as a binary classification task. Our second use case is derived from the biomedical domain where we use human gene expression data to predict cell response to a series of infections (Influenza, E. coli and control) as a multi-class classification task. Finally, our third use case focuses on environmental data where we predict soil pH (a regression task) from normalized soil microbiome species abundance data. In each instance we use algorithms including Random Forest (RF), XGBoost (XGB), AdaBoost, K-nearest neighbours (KNN), LightGBM (LGBM) and a neural network (Auto Keras) followed by XAI to assess the biological validity of the ML model i.e., to check if key predictors align with current biological know-how. We show that the XAI component of the workflow can act as both a ML model validator in this way, but also as a hypothesis generator to rank key omic features that are driving the predictions for a target or phenotype of interest since these top features are candidates for further analysis or experimental testing.

## Results and Discussion

### Binary classification using AutoXAI4Omics: a case study in plant genomics

ML and high throughput phenotyping within plant sciences can reduce laborious manual work and cost while improving the efficiency of data collection e.g., via automated image collection and analysis [28]. ML has also been used to improve the time-lines for plant breeding [29, 30]. Omics datasets can be utilised to select genes that underpin traits of interest to enable their incorporation into breeding programs [31]. ML can fine tune the selection of SNPs or QTLs from genomic datasets within genome-wide association studies and genomic selection. New ML tools are being continually developed for understanding traits of interest in plants, for investigating gene expression [17] and for integrating multi-omic datasets to predict phenotypes [32, 33].

We downloaded a genotyping matrix of global GBS (genotype-by-sequencing) information plus related phenotype data for 1000 core Barley (*Hordeum vulgare L.*) accessions published in [34]. In this data set Barley accessions are categorised as two-rowed or six-rowed based on their lateral spikelets and floret sizes. Six-rowed Barley has three spikelets all of which are fertile and can produce grain. While two-rowed Barley has three spikelets, but only the central spikelet is fertile [35]. We converted phenotypic data into binary format; two-rowed (0) or six-rowed (1) and converted the features or SNPs into numerical format for usage in AutoXAI4Omics; homozygous for reference allele (2), heterozygous (1), homozygous for alternative allele (0) and NNs (3). The data set thus included 957 Barley accessions (after removing those with intermediate phenotypes or missing data) and 37,953 features (SNPs) which were used to train ML models to perform classification for row-number.

AutoXAI4Omics was run for this data set using classification mode, a random search, f1-scoring and a train:test split ratio of 80:20. The config file that was used to run this analysis is provided as File-S1.json and the associated data/metadata files to run this config as File-S2.csv and File-S3.csv respectively. Figure 2 summarises our results, where we also used feature selection for 1000 features, a number determined manually based on the feature selection accuracy in Figure 2-c. Figure 2-a shows the performance of a range of hyper-tuned ML models for the prediction of the target during cross validation. Several models perform well, though no two are identical. Our encoded algorithm within AutoXAI4Omics suggests the “best” or “recommended” model for the user to focus on (here XGBoost)-elements of this decision are visible in Figure 2-a where XGBoost is generating a high accuracy coupled to a low performance variation across the folds on cross validation. What is not shown in Figure 2-a is additional information (considered by AutoXAI4Omics) regarding minimal overfitting observed when comparing training and test data performance (f1-score) and balanced performance across the predicted classes (i.e., performance was most balanced for XGBoost). Figure 2-b depicts the confusion matrix and ROC curve (Figure 2-d) for the XGBoost model, its F1-scores were 1.0 on the training data and 0.99 on the test data (these values and others are reported as a results table for the user; Table S1).

**Figure 2:**
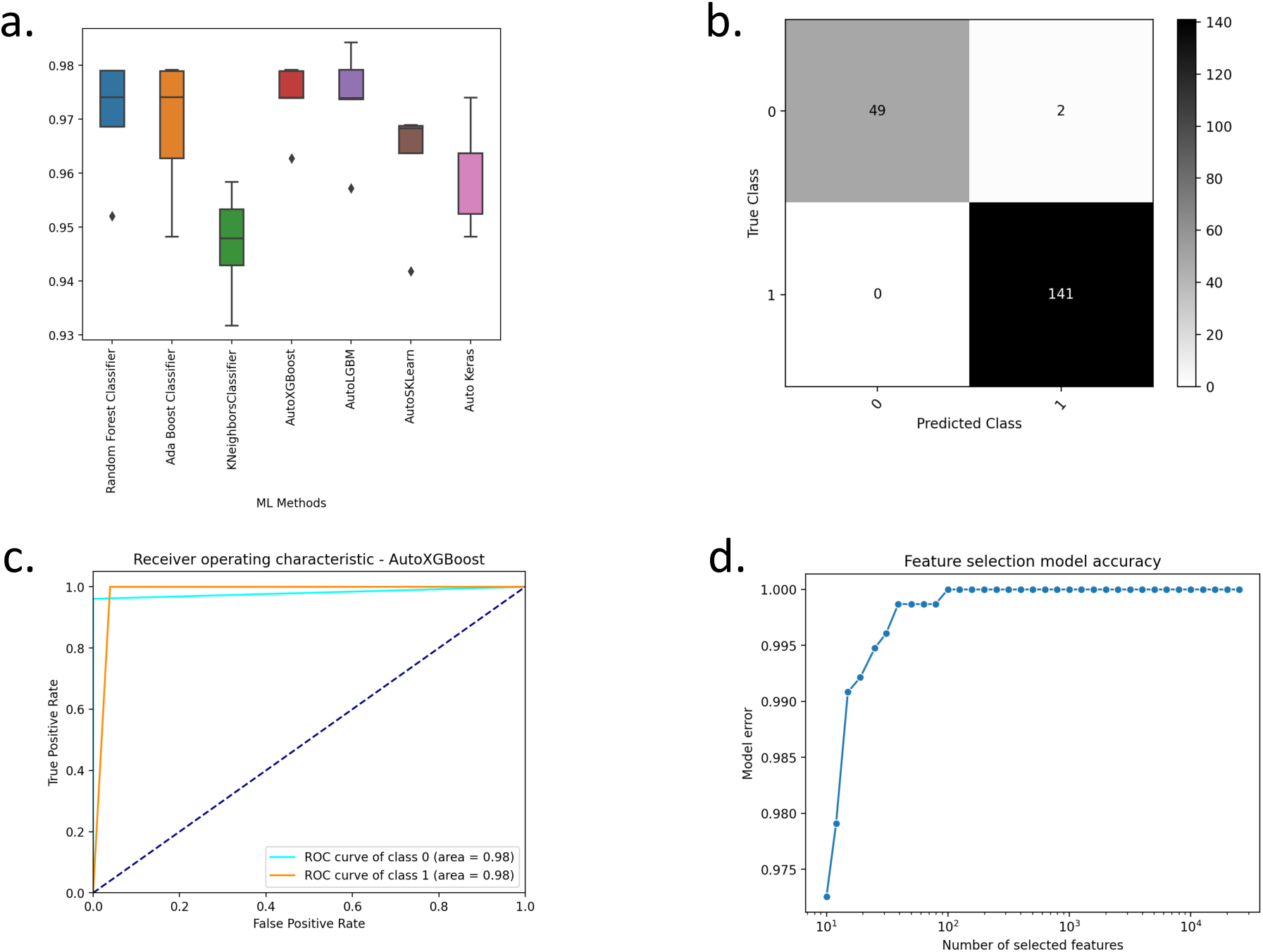
Binary classification using AutoXAI4Omics: a case study in plant genomics. Figure summarises the performance of the AutoXAI4Omics supervised ML workflow to predict either two-rowed (0) or six-rowed barley (1). (a) Box plots displaying the f1-score during cross validation (b) Confusion matrix for the best performing ML model as identified by AutoXAI4Omics (XGBoost), (c) ROC curve for the best performing model (XGBoost), (d) Feature Selection Accuracy curve as identified by AutoXAI4Omics using a default Random Forest classifier.

Figure 3 summarises the XAI results for the Barley binary classification. Highlighting the top 15 features and their relative importance’s for the classification via SHAP. This allows us to interpret the results and validate the ML by linking predictive features to biological processes. For example, the two most predictive SNPs for row number in Barley from AutoXAI4Omics are within chromosome 2 at position 2:651372029 and 2:651378029 in order of their relative importance. We searched for genes within a 20kb region around these SNPs utilising GrainGenes [36]. Two high confidence genes were found within this region based on the Barley cv. Morex V3 annotation [37]; HORVU.MOREX.r3.2HG0211880 (Exostosin family protein) and HORVU.MOREX.r3.2HG0211900 (Phytol kinase 1). HORVU.MOREX.r3.2HG0211900 has been associated with the differentiation of two-rowed and six-rowed Barley in prior research [38], utilising Barlex [39]. This gene may also be linked to the loci *six-rowed spike vrs*, *hexastichon (hex-v)* or *intermedium spike (int)* which are heavily discussed in the literature due to their association with row number in Barley [40, 35, 41, 42, 43]. Further evidence to support that the genes identified are linked to row number and spike development in Barley is from expression data, which suggests their expression during the seed development and within spike meristems [44, 45]. Finally, orthologues in Wheat (*Triticum aestivum)* with over a 90% sequence match to the genes from Barley were found using ensembl [46]. The orthologues were then investigated further utilising KnetMiner [47] which associated them to pathways within root and flower meristems and for seed development and germination.

**Figure 3:**
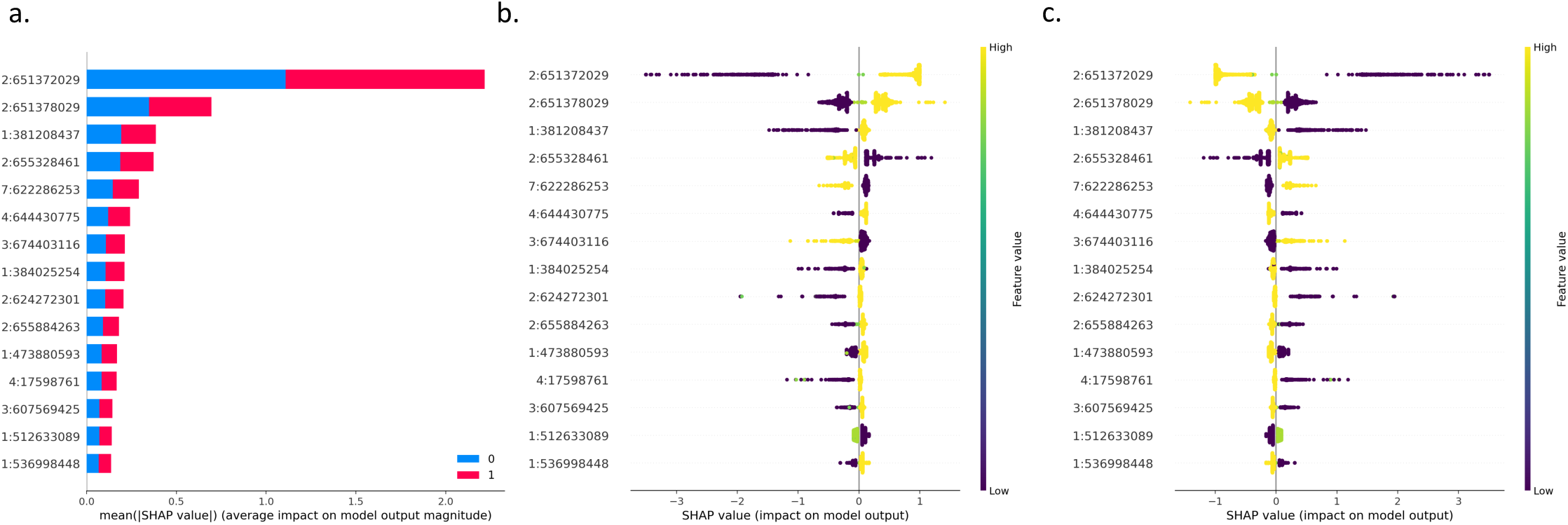
Binary classification using AutoXAI4Omics: a case study in plant genomics. Figure summarises the XAI output relating to the best performing model from AutoXAI4Omics(XGBoost) to predict either two-rowed (0) or six-rowed barley (1). The y-axis displays the results of the XAI analysis which ranks the relative importance or impact of the input features (SNPs represented as chromosome:position) on the best ML model (XGBoost) – features are ranked from high impact to low from top to bottom of the y-axis. (a) Bar plot summarising the top 15 feature values selected for the best model (XGBoost) relative to their average impact on the model predictive output, (b, c) SHAP summary dot plots displaying the top 15 features (y-axis) against the Shapley value (x-axis) and the colour represents the feature values (SNPs encoded as 0, 1 or 2). A positive SHAP value for a feature represents a positive association with that feature value and the sample being assigned to the specific class being visualised in the plot; (b) Explanation for six-rowed, (c) Explanation for two-rowed Barley.

### Multi-class classification using AutoXAI4Omics: a case study for human RNA-seq

We downloaded and used the Series Matrix text file for study GSE53165 that is part of the NCBI Gene Expression Omnibus. This data set was generated as part of a prior publication [48] and is related to the variation in individuals’ responses to complex diseases in humans. The authors generated transcriptional profiles from dendritic cells (DCs), derived from the peripheral blood of healthy individuals that were subjected to different states. Here we compare three of these states; resting (unstim), E. coli lipopolysaccharide stimulated (LPS) and influenza-infected (dNS1). We used this data set, encompassing 2009 samples (that we reduced to 167 by removing serial replicates) and 414 features (genes with expression information), to train a series of ML models to perform multi-class classification. The targets to predict were the three treatments unstim, LPS and dNS1. The config file used to run this analysis in AutoXAI4Omics is attached as File-S4.json and the associated data/metadata files to run this config as File-S5.csv and File-S6.csv respectively. Most notably for this analysis, we used classification mode, a random search, f1-scoring, automated feature selection and automated filtering for samples more than 5 standard deviations away from the mean and those with adjusted gene expression counts of 1 or less in 10 or less samples. Figure 4 summarises the results from the AutoXAI4Omics analysis, with Figure 4-a showing the performance across all models (test data), and Figure 4-b-d focusing on our best performing model as identified by AutoXAI4Omics (Random Forest) where the confusion matrix (Figure 4-b) and ROC curves (Figure 4-d) highlight robust performance across all classes. The weighted F1-scores across the three classes for our best model were 0.99 on the training data and 0.97 on the held-out test data (Table S2).

**Figure 4:**
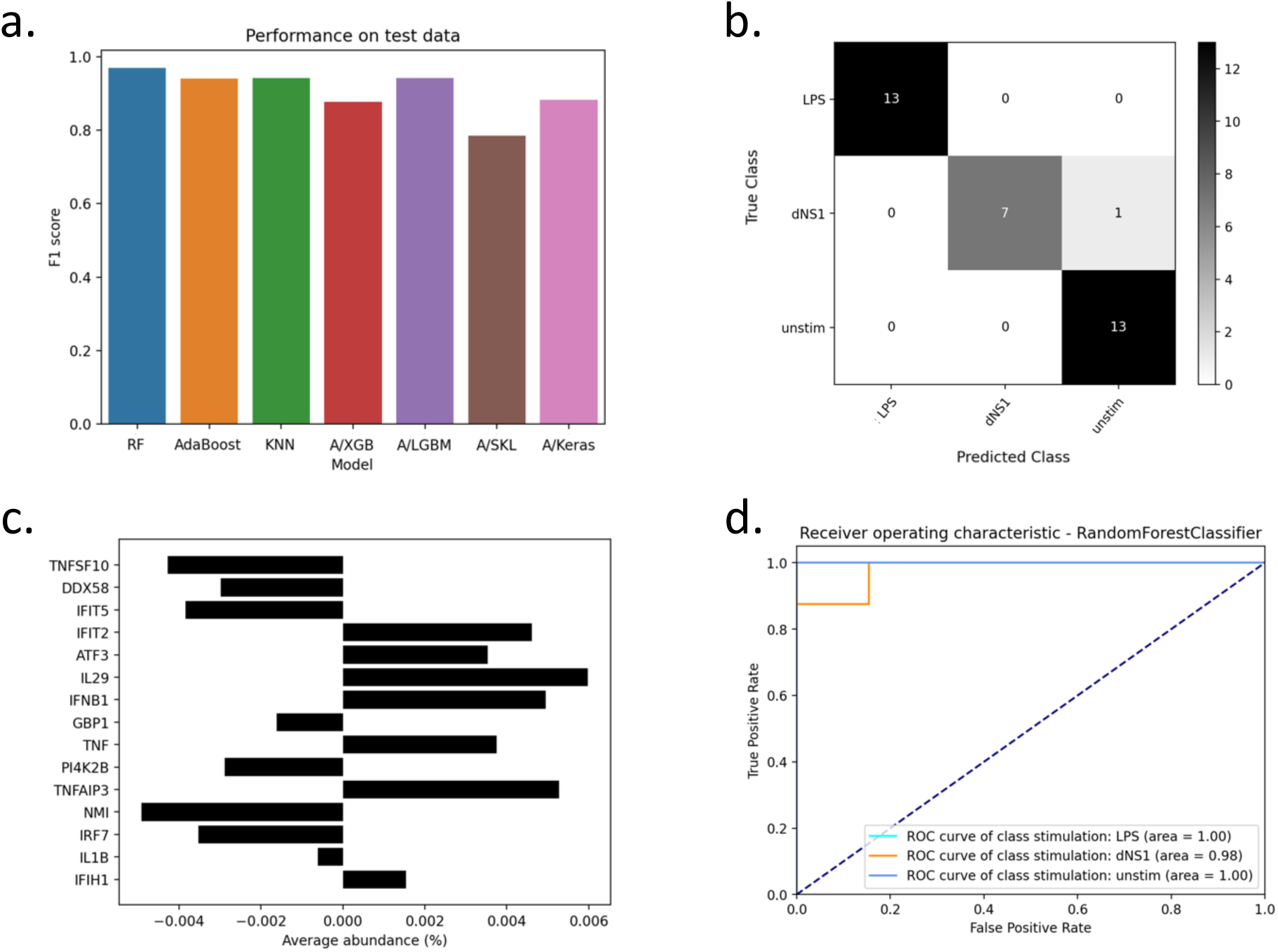
Multi-class classification using AutoXAI4Omics: a case study with human RNA-seq data. Figure summarises the performance of the AutoXAI4Omics supervised ML workflow to predict three classes unstim, LPS and dNS1. (a) Bar charts displaying the f1-score on the held-out test dataset (b) Confusion matrix for the best performing ML model as identified by AutoXAI4Omics (Random Forest), (c) Bar chart to display results of feature importance analysis to rank the relative importance of the input features prior to ML model development (y-axis), the x-axis depicts the average abundance of the related features across the input sample set (average gene expression counts) (d) ROC curve for the best performing ML model as identified by AutoXAI4Omics (Random forest).

Figure 4-c gives insight into the relative importance’s of the input features to the model, but the advantage of our XAI workflow is a more comprehensive interpretation of the generated models via SHAP. Figure 5 shows the SHAP output for our best performing ML model (Random Forest) for each of the three classes. We often see features having a mirror image role in different classes (as per binary classification) e.g., for the most predictive gene (dNS1 class) Interleukin-29 (IL29), a higher abundance positively associates with a sample being linked to the dNS1 class, while a lower abundance positively associates with a sample being linked to the LPS class. Furthermore, we advocate using these XAI outputs to sanity check the biological validity of the ML model i.e., to check if key predictors align with current biological know-how. In support of this, the most predictive gene for class dNS1 (also the fourth most predictive for class LPS), was IL29. IL29 is a cytokine that has been linked to antiviral activity, antibacterial activity, antiproliferative activity and in vivo antitumour activity [49], and higher IL29 expression has been linked to influenza infection i.e., the dNS1 class (which is the pattern reflected in Figure 5-b). Conversely, for LPS induction of IL29, previous work noted high expression of IL29 in DCs after initial LPS stimulation (0-2 hours) and then a steep drop-off in IL29 expression after further LPS exposure (2-18 hours) [50]; since the cell’s used in our ML analysis were exposed to LPS for more than 2 hours (up to 5 hours) this aligns with the low-mid expression of IL29 driving a prediction of class LPS (seen in Figure 5-a). Fittingly IL29 is not present in the most predictive features for the unstim class where no infective agent is present.

**Figure 5:**
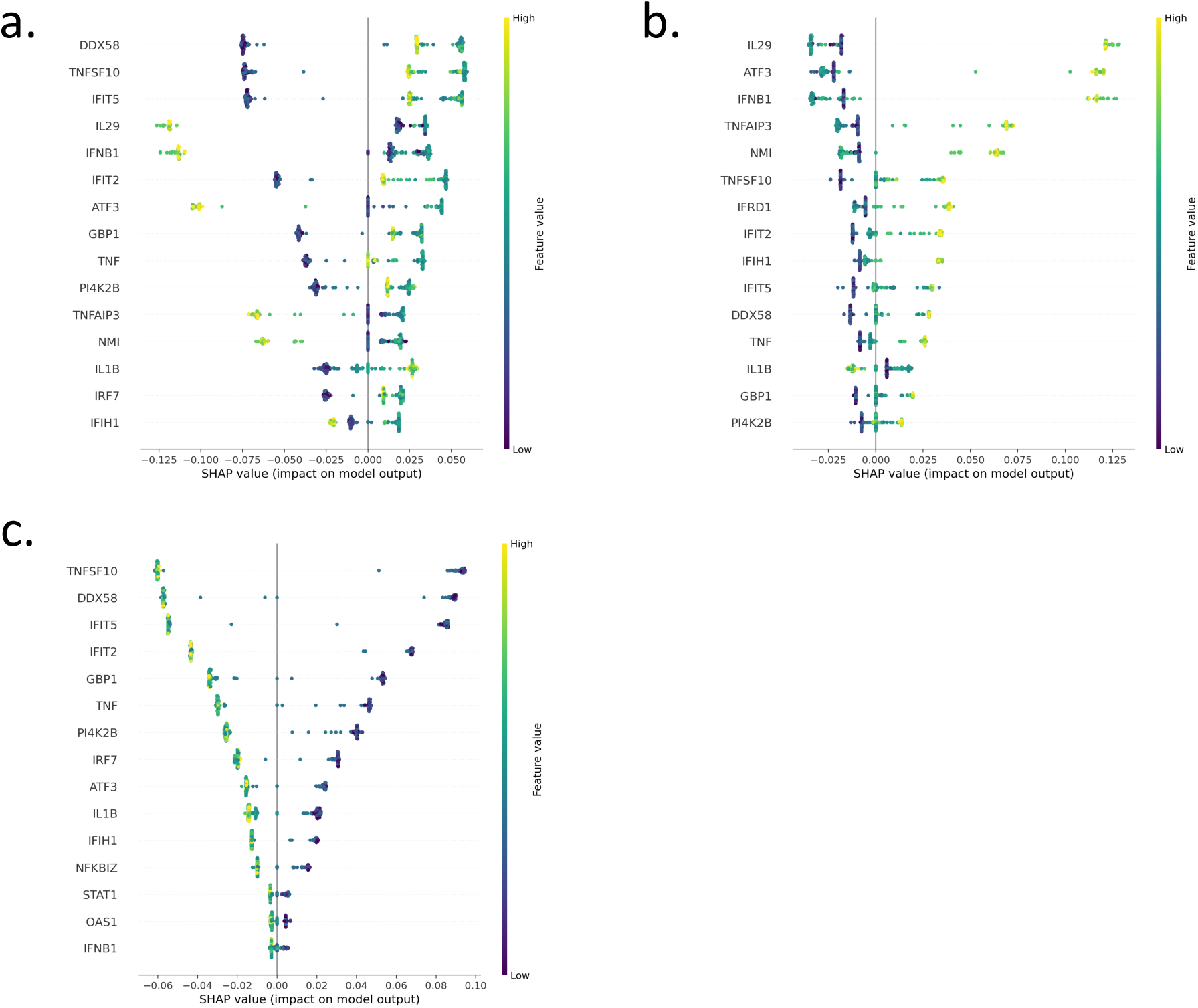
Multi-class classification using AutoXAI4Omics: a case study with human RNA-seq data. Figure summarises the XAI output relating to the best ML model generated by AutoXAI4Omics (Random Forest) to predict three classes unstim, LPS, and dNS1. (a) Global explanation for class LPS, (b) Global explanation for class dNS1, (c) Global explanation for class unstim.

Many other predictive genes align with common sense as to what might be relevant for response to disease or stress e.g., Activating transcription factor 3 (ATF3) a stress induced transcription factor where high levels of this gene make a sample more likely to be assigned to the dNS1 class, mid-expression levels of the gene are associated with the LPS class and low levels with the unstim class. Also, Guanylate-binding protein 1 (GBP1), that plays important roles in innate immunity against a diverse range of bacterial, viral and protozoan pathogens and follows broadly the same predictive profile as ATF3 i.e., high levels make a sample more likely to be assigned to the dNS1 class, mid-expression levels are associated with the LPS class and low levels with the unstim class.

### Regression using AutoXAI4Omics: a case study for environmental microbiome data

We used a bioinformatics workflow (see Methods) to generate species abundance information for 189 environmental microbiome samples that were part of a study presented by Bahram et al. [51] comprising whole shotgun metagenomes from samples of topsoil collected from representative terrestrial regions and biomes across the world. Based on sample geo-locations, we gathered information for each sample regarding the pH of the soil at the sampled depth (0 to 5 cm) from SoilGrids [52]. We used this data set, encompassing 189 samples (that we reduced to 173 by removing low quality samples) and 91,837 features (species with normalised abundance information), to train a series of ML models to predict pH as a regression task. The config file used to run this analysis in AutoXAI4Omics is attached as File-S7.json and the associated data/metadata files to run this config as File-S8.csv and File-S9.csv respectively. To run this analysis, most notably, we used regression mode, a grid search, Mean Absolute Error (MAE) scoring, automated feature selection and automated filtering for samples more than 5 standard deviations away from the mean and those with abundances of 5 or less in 10 or less samples.

Figure 6 summarises the results from the AutoXAI4Omics analysis, with Figure 6-a showing the performance across all models on the held-out test data, Figure 6-b showing this performance on cross validation, and Figure 6-c-d focusing on our best performing model as identified by AutoXAI4Omics (Random Forest), where the correlation plot (Figure 6-c) and joint plot (Figure 6d) highlight the high degree of correlation between true and predicted values. The MAEs for our best model were 0.02 on the training data and 0.31 on the held-out test data. AutoXAI4Omics also allows us to output different evaluation metrics and we found the metrics Mean Absolute Percentage Error (MAPE) and R2 can also be useful to assess performance. For this analysis there was a MAPE of only 5.7% on the held-out test data a relatively high R2 of 0.64.

**Figure 6:**
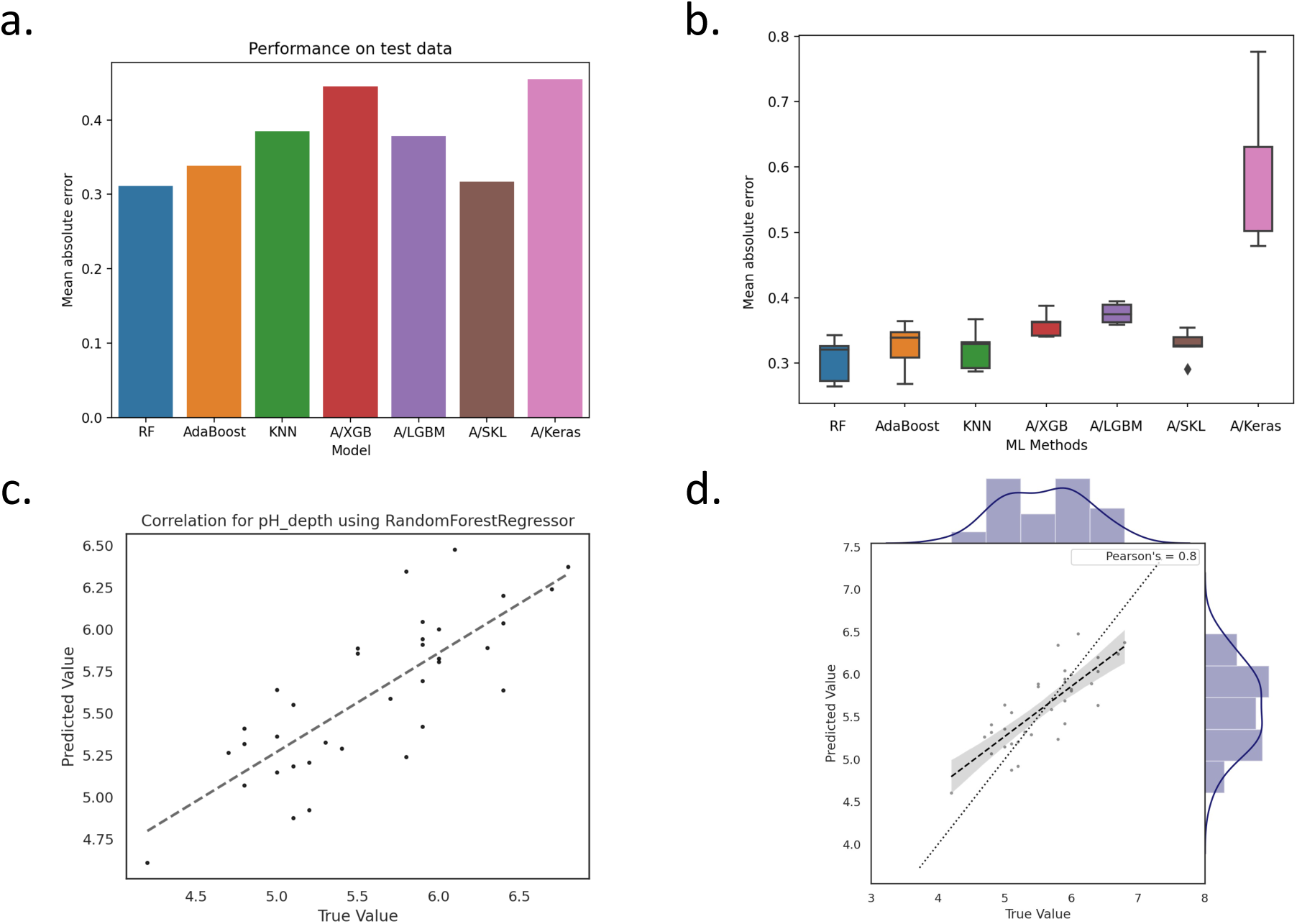
Regression using AutoXAI4Omics: a case study with environmental microbiome data. Figure summarises the performance of the AutoXAI4Omics supervised ML workflow to predict soil pH from the microbiome. (a) Bar charts displaying the MAE on the held-out test data set across a range of ML models (b) Box plots to show the MAE on cross validation across a range of models, (c) Correlation and (d) Joint plot to compare predicted values (y-axis) with true values (x-axis) for the best performing ML model as identified by AutoXAI4Omics (Random forest). Diagonal line is also shown, dotted line.

Figure 7 shows the SHAP output of the top 15 most predictive species for our best performing ML model (Random Forest) for soil pH. Notably, these species include several acidophiles or acid-linked species e.g., Acidocella sp. MX-AZ02, Ralstonia virus RSL1 and an Acidobacteria bacterium. As one might expect these species show a positive association with the prediction of a low pH i.e. higher species abundances tend to associate with lower or negative SHAP values that lower the models predicted pH for a sample. Additionally, the most predictive species Candidatus Nitrosotalea sp.FS has known associations with acidic soil that fit both its appearance as a predictor but also the directionality of its effect (i.e., higher abundance predictive of a lower pH). Moreover, species within its associated genus, Candidatus Nitrosotalea, are reportedly ammonia oxidisers abundant in acidic soil environments and obligate acidophiles, growing optimally around pH 5.0 which directly supports our observed importance and trends [53].

**Figure 7:**
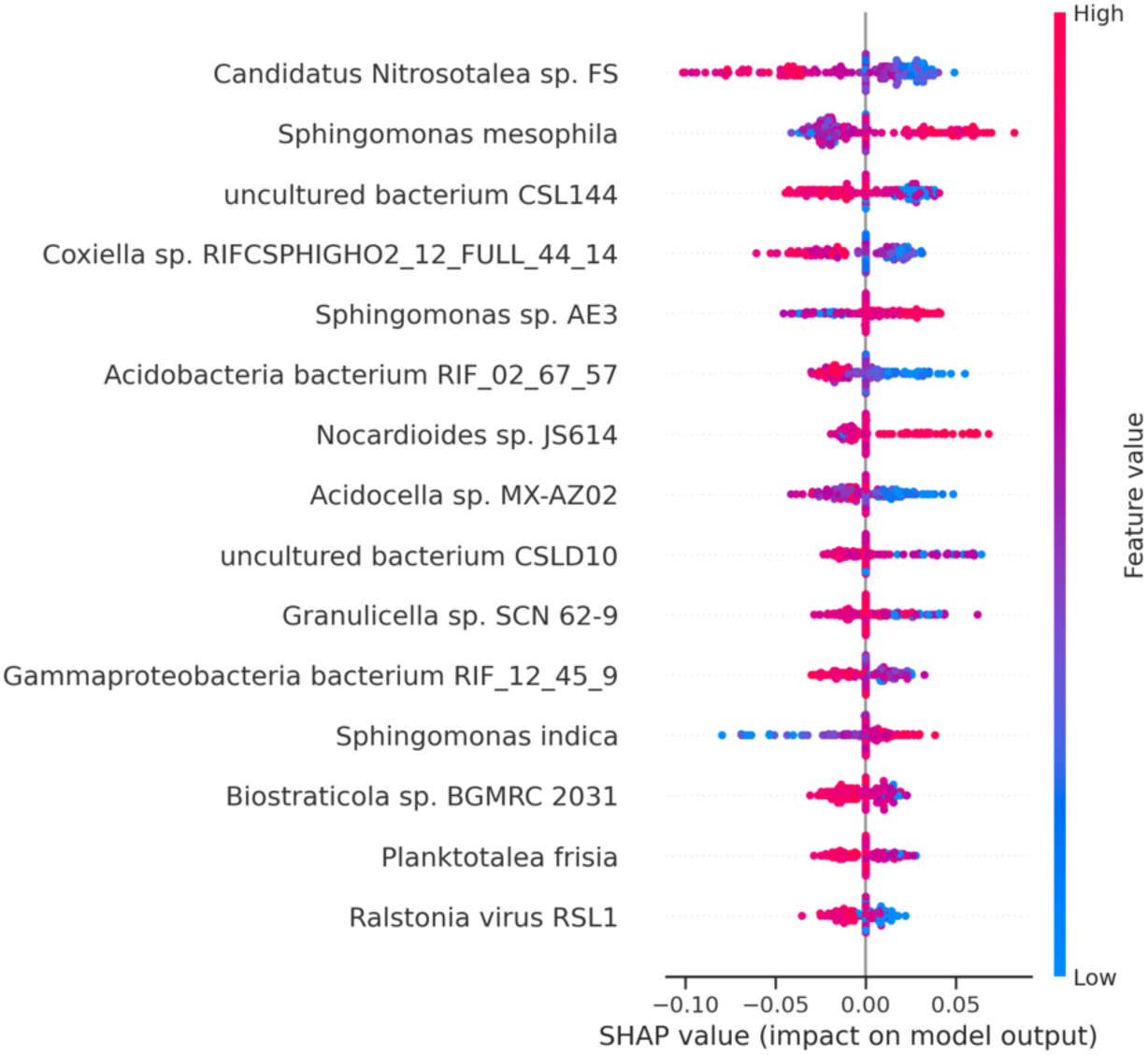
Regression using AutoXAI4Omics: a case study with human RNA-seq data. Figure summarises the XAI output relating to the best ML model generated by AutoXAI4Omics (Random Forest) to predict soil pH. Global explanation is shown where a low feature value (blue) indicates a low species abundance, and a higher feature value (red) indicates a higher species abundance). If a higher feature value correlates with a higher SHAP value (right hand side of the plot axis) then a higher species abundance tends to predict a higher pH value. Conversely, if a higher feature value correlates with a lower SHAP value (left hand side of the plot axis) then a higher species abundance tends to predict a lower pH value. Some full names have been shortened as follows: Gammaproteobacteria bacterium RIFCSPHIGHO2 12 FULL 45 9 shortened to Gammaproteobacteria bacterium RIF 12 45 9 and Acidobacteria bacterium RIFCSPHIGHO2 02 FULL 67 57 shortened to Acidobacteria bacterium RIF 02 67 57.

### Conclusions

In this study we present our open-source end-to-end explainable ML tool, AutoXAI4Omics. We apply it to three use cases, and we provide a series of examples (including related data and configuration files) to showcase the utility and performance of the tool. Our use cases cover a wide range of omic data types (genomics, transcriptomics, microbiome), application domains (plant science, environment and human disease), tasks (multi-class/binary classification, regression) and pre-processing options (automated feature selection, user defined feature selection, feature/sample filtering). We also focus on the advantage offered by the inclusion of XAI approaches alongside feature importance techniques. In doing so we highlight the wide applicability of our tool to a wide range of research areas to help scientists answer a variety of life sciences questions. AutoXAI4Omics can be used by domain scientists that are not ML experts to quickly perform robust and trustworthy interpretable ML analysis and therefore focus on the insights generated by the tool, rather than in building and fine tuning ML models, hence accelerating scientific discovery.

## Methods

### AutoXAI4Omics workflow

AutoXAI4Omics has four modes that can be invoked, which are:

- *training* – Train, tune and evaluate several ML algorithms to perform classification or regression tasks on a given dataset as specified in the configuration file.
- *plotting* - Create (or re-create) plots and visualisations for a trained and optimised model or a set of models as specified in the config.
- *holdout* - Apply the trained and tuned model(s) to a holdout separate dataset and evaluate their predictive performance on it.
- *prediction* – Makes predictions on new, unlabelled data, using the best trained model and get its explanations.

The flow that we describe below includes those steps that all belong to and run in the training mode. The other modes invoke the corresponding stages that are required for their individual tasks, e.g., plotting only or predicting only on holdout dataset or new unlabelled data points.

### Load data *&* data specific processing

The first step AutoXAI4Omics performs is to load the data into the system. At this point if the user chooses, omic-specific pre-processing is performed on the data. This is an optional step and is only done if the user sets the corresponding field in the configuration file, data_type. We enable this step to be optional as there are many ways that this pre-processing can be done, with users possibly having their own preferred method. If a user has already pre-processed their data, then they may delete the omic specific entry from the config and this will stop any pre-processing from happening. Also AutoXAI4Omics can accept and work with non-omic data, if a user wishes, and as before, the user needs only remove the omic-specific pre-processing field from the config. AutoXAI4Omics includes pre-processing modules for gene expression data, microbiome data, metabolomic and tabular numerical data.

The gene expression pre-processing is designed to accept RNA-seq read counts and microarray expression levels and is activated when the user specifies in the config data type=”gene expression”. The user needs to also specify the type of their expression data e.g., ’FPKM’, ’RPKM’, ’TMM’, ’TPM’, ’Log2FC’, ’COUNTS’, ’OTHER’, where ‘COUNTS’ invokes conversion of raw count data to normalised trimmed mean of M-values (TMM) values. Other additional pre-processing capabilities that are popular for those using gene expression data, have been added to allow filtering of unwanted or potentially problematic genes or samples. Firstly, the function “filter sample” removes samples if the number of genes with coverage is more than X standard deviations from the mean across all samples i.e., the extremes of very sparse or high coverage samples. Secondly, the function “filter genes” removes genes unless they have a gene expression value over X in Y or more samples i.e., removing genes that never appear with sufficient expression across the sample and may be noise.

The omic-specific pre-processing also includes several functions specifically for microbiome data. These functions are available when the user specifies data type=”microbiome” in the configuration file. Microbiome-specific pre-processing capabilities include optional collapsing of taxonomy to kingdom; ’k’, phylum; ’p’, class; ’c’, order’ ’o’, family; ’f’, genus; ’g’, or species; ’s’ levels, which is invoked by the user specifying the taxonomic rank letter in the “collapse tax” option in the configuration file. Two read-based filtering options are available; “min reads”, which will remove samples with fewer reads than specified and “norm reads”, which will rescale all samples to sum the number of reads specified. Both “min reads” and “norm reads” have default values of 1000, which will be invoked if data type=”microbiome” is selected and the user does not change the default values in the configuration file. Taxon-based filtering is available with functions “filter abundance”, which will remove taxonomic features with a total count less than the specified number across all samples, and “filter prevalence”, which will remove taxonomic features which occur in less than the specified proportion of samples. The default value for “filter abundance” is 10. For “filter prevalence” the default value is 0.01, which represents the decimal percentage of the samples, so using the default value here, any taxonomic features present in less than 1% of samples would be removed. Metadata-based filtering is also supported for microbiome data. The function “filter microbiome samples” allows for removal of samples based on metadata categories. This function requires a dictionary (or list of dictionaries), where the key represents the metadata column (i.e. ’COUNTRY’) and the value represents the metadata category in that column to be removed (i.e. ’UK’). The default value for this function is ’null’, invoking no metadata-based filtering. Two final functions are available which can manipulate the ’target’ metadata category specifically. These include “remove classes”, which takes in a list of class labels to be removed from the ’target’ column, and “merge classes”, which allows for merging of target classes by taking in a dictionary in the format ’{”X”: [”A”, ”B”]}’, which would convert class labels ’A’ and ’B’ to ’X’. Both “remove classes” and “merge classes” have default values of ’null’, invoking no manipulation of the ’target’ column.

The omic-specific pre-processing also supports data types “tabular” and “metabolomic” and these have associated filtering options, similar to the gene expression data, for i) removing samples if no of measures with measurements is more than X std from the mean across all samples, and ii) for removing measures unless they have a value over X in Y or more samples.

AutoXAI4Omics only loads and pre-processes one data type per run. For now, if a user has a data set that has non-omic and omic features then they can do any pre-processing first, outside of the tool themselves, and then give it to AutoXAI4Omics as a ”non-omic” data set. This ability to intergrate multiple, and different, omic types automatically is a feature that we plan to bring to the tool in future. One other feature that we plan to add later is the automated encoding of any categorical features. For now if the user has any categorical features, then they must manually convert them into one-hot-encoded columns first.

### Split data

Once the previous step has been completed the data is split into train-test sets as characterised by the setting in the config that has been presented by the user. If not presented, then the testing size will default to 20%. The exact implementation is done by scikit learns’ *train test split* [54]. The split will depend on the random seed that has been set, which also impacts the initial model weights and performance. This can be set in the config provided by the user, but if not provided will be set to a default value.

### Standardise data

Next the tool standardises the columns of the training data, for which we use scikit-learns’ *QuantileTransformer* [54]. Ideally when training AI models data should be normally distributed, but in real world situations this is rarely the case and even standard normalisation is not completely immune to skewed datasets. This is the reason why we choose to use the *QuantileTransformer* as it is more robust to skewed data.

### Feature Selection (optional)

Feature selection then follows this step, and is completely optional, if the user does not wish feature selection to take place then they need only remove the corresponding entry from the config. However, if the user does wish to perform feature selection on the training data then there are several options that they can customise/set. The first step that occurs is variance thresholding, (*variance removal* [54]), to remove constant or near-constant columns. The value for this can be set by the user but if not provided will default to 0. Once this is done the tool can either search for a specified number of best features or it can automatically find the best number of features.

In the case of finding a specified number of best features then the tool will find them using the setting given by the user, if not present it will default to using *SelectKBest* method with either the *f classif* or *f regression* metric as implemented by scikit learn [54]. *SelectKBest* works by taking the scoring metric provided to it and select the K best scoring features. If using *f classif* this is the ANOVA F-value of the column in relation to the targets and *f regression* works by running univariate linear regression tests with respect to the target.

If the user then chooses to let the tool automatically find the best number of features they can choose to set the minimum and maximum number of features the tool will search between. For efficiency the tool creates several potential candidates that are logarithmically spaced between these two values. Then for each candidate number a set of best feature features are found and a default model is trained and evaluated on it. Once all candidates have been scored we then use our own unique method to select the best candidate number of features to proceed through to the next step. To select the value for K we look for a set of 3 consecutive values that are the best performing and the most stable, once we have that then we choose the largest k in that set as the candidate to use. By stable we mean that the set has a low standard deviation and we wanted to factor this in is to make sure that our selection is robust. Hypothetically, if an optimum K that has terribly performing neighbours, making a ”V” shape then this k should be treated with an abundance of caution as the addition or removal of a feature causes a drastic chance and could have been achieved because of any number of reasons.

### Class balancing (optional, problem specific)

Class balancing is only available when performing classification problems and is also optional. If a user does not want to implement, then they just need to set the corresponding variable in the config. If however, they do wish to perform class balancing then there is the option to perform both Over- sampling and Under-sampling, implemented by imblearn [55].

### Train *&* evaluate models

Now the training of the models can begin. Models available to train include those from scikit learn[54], Auto-scikitlearn[56], xgboost[57], lightgbm[58] and autokeras[59]. These can be trained with most standard metrics that are available, such as: Accuracy, F1 score and Precision for classification tasks; Mean squared error, Mean absolute percentage error and R^2^ for regression tasks. In addition, these models can also be hypertuned on the training data with random or grid search, or without if so desired. A full explanation of all the possible combinations can be found in the code repository. The exact models that are trained are controlled by a corresponding entry in the config the user provides along with what metric they want to fit according to and what hyper-tuning to use.

### Plotting

Once training is complete, if the user desires plots can be produced illustrating the model performance and explaining how the models work. These plots are optional and the exact ones produced are controlled by a corresponding list in the users config, meaning it can be skipped if set to empty.

But if plots are desired, these are produced using a combination of seaborn[60], matplotlib[61] and scikit learn[54]. The full range of plots are described in our code repository but examples are bar & box plots of model performances plus roc curves and confusion matrices if performing a classification task. Many of these are shown in the Results and Discussion section of this paper. For the explainability aspect, the two libraries we utilise are eli5, for permutation importance, and shap[21], for shap value bar and dot plots - again examples of these can be seen in Figure 3, 4, 5, 6 and 7. The user can also define and control the number of feature that are calculate/plotted via the config file and also specify what data the shap values are to be calculated based on i.e. the training set, test set or both.

### Select best model

The last step in the process is to select the best model out of those that were trained during the run of AutoXAI4Omics. This selection is done with respect to the metric that the user sets for the models to optimise. For both problem types, classification and regression, the training error and test error are collected and plotted. In the regression problem, the point that is closest to the y=x and the origin (0,0) is selected as the best model. For classification problems, it is the same but we switch the origin for the unit point (1,1) as in classification problems the higher performance value the better and values are bounded above by 1. Sometimes this may result in ties, in which case AutoXAI4Omics will pick one but make a point of noting this in their own text file. Once the best model is selected all of the related content is copied to a dedicated folder ”best model”.

### Datasets

In the Results and Discussion section we showcase running AutoXAI4Omics with three types of data including genomics, transcriptomics and microbiome abundance data. For transcriptomic data this tends to be numerical, and so by nature is compatible with most ML algorithms, but we recommend using normalisations to optimise between sample comparisons e.g., such as TMM that we include in AutoXAI4Omics pre-processing. This numerical nature also applies to microbiome data where users can input normalised or rarefied counts or relative abundance data. Here we also recommend testing centered log-ratio (CLR) transformation that removes compositional artifacts. In contrast, genomic data such as SNPs, can commonly be encountered as a series of alleles e.g., for diploid organisms we might see homozygous reference allele calls, heterozygous calls and homozygous alternate allele calls. For such data we recommend converting these calls to numerical representations. We have found increasing representative numbers from 0-2 as the presence of an alternate allele increases i.e., moving from homozygous reference (0) to heterozygous (1) and then homozygous alternate calls (2), can be an intuitive way to represent this type of data, allowing more meaningful interpretation downstream e.g., to assess as the alternate allele increases what is the effect on the predicted target variable.

Preparation of our three example types of data (genomic, transcriptomic and microbiome) for usage with AutoXAI4Omics is detailed in the main text. For the genomic and transcriptomic data processed data was available from public sources that we state. However, we generated the microbiome abundance data from raw sequencing reads that were available as part of the study by Bahram et al. [51] at the European Bioinformatics Institute -Sequence Read Archive database under PRJEB24121. We used a bioinformatic analysis workflow for the taxonomic annotation of paired end reads. In our workflow, using Trimmomatic v2.9 [63], reads were trimmed of adapter and if the average quality across a 4bp window was less than 15, reads below 40bp were dropped from the analysis. Reads were then used as input into DIAMOND v2.0.11 [64] for read alignment in blastx mode using “-b 25 -k 5 – index-chunks 4 –min-score 50 and –max-target-seqs 2” options. For these DIAMOND aligned reads the suite of tools from the MEGAN community edition v6.21.12 [65] were used to run a last common ancestor (LCA) analysis to allow binning of read pairs by taxon. MEGAN outputs were then processed to extract and compute species level abundance (read counts).

## Supporting information

File-S1.json

File-S2.csv

File-S3.csv

File-S4.json

File-S5.csv

File-S6.csv

File-S7.json

File-S8.csv

File-S9.csv

## Declarations

### Consent for publication

All authors have consented to publication of the manuscript. No additional consent required.

### Availability of data and materials

The experimental datasets that were used in this study are all publicly available and details of how to access them can be found in the main text at the first mention of the dataset. The processed datasets used directly in AutoXAI4Omics are provided with this manuscript as files S1-9.

### Authors’ contributions

All authors contributed code and/or conceptualisation of AutoXAI4Omics functionality. LJG, KD-J and JS performed manuscript writing, APC and LJG performed manuscript review and editing.

## Acknowledgements

This work was supported by the Hartree National Centre for Digital Innovation (HNCDI), a collaboration between STFC and IBM. KD-J PhD was supported by UKRI-BBSRC through the Norwich Research Park Doctoral Training programme (#2578607).

