## Supplementary information - Tables S1-2 for "AutoXAI4Omics: an Automated Explainable AI tool for Omics and tabular data"

Supplementary Tables S1-S2

Supplementary Data files S1-S9 (formats .json, .csv)

**Supplementary Tables**

**Table S1. Output performance metrics for ML models generated from AutoXAI4Omics for example task: *Binary classification in plant genomics (Barley).*** This is the default table generated as output in folder “Results” file name: scores__performance_results_testset.csv. Train refers to the training dataset for each ML model and Test refers to the held-out test dataset used for model evaluation.

| **model** | **accuracy_score_Train** | **accuracy_score_Test** | **f1_score_Train** | **f1_score_Test** | **f1_score_PerClass_Train** | **f1_score_PerClass_Test** |
| --- | --- | --- | --- | --- | --- | --- |
| **RandomForestClassifier** | 0.9973856209150330 | 0.9791666666666670 | 0.9973856209150330 | 0.9791666666666670 | [0.99509804 0.99821747] | [0.96078431 0.9858156 ] |
| **AdaBoostClassifier** | 1.0 | 0.9791666666666670 | 1.0 | 0.9790316901408450 | [1. 1.] | [0.96       0.98591549] |
| **KNeighborsClassifier** | 0.9790849673202610 | 0.9739583333333330 | 0.9793283064360210 | 0.9740384117057670 | [0.96226415 0.98553345] | [0.95145631 0.98220641] |
| **AutoXGBoost** | 1.0 | 0.9895833333333330 | 1.0 | 0.9895158450704230 | [1. 1.] | [0.98       0.99295775] |
| **AutoLGBM** | 0.9816993464052290 | 0.9791666666666670 | 0.9816114463145270 | 0.9790316901408450 | [0.96517413 0.98758865] | [0.96       0.98591549] |
| **AutoSKLearn** | 0.9816993464052290 | 0.9635416666666670 | 0.9816993464052290 | 0.9634255938844770 | [0.96568627 0.98752228] | [0.93069307 0.97526502] |
| **AutoKeras** | 1.0 | 0.9791666666666670 | 1.0 | 0.9792925824175830 | [1. 1.] | [0.96153846 0.98571429] |

**Table S2. Output performance metrics for ML models generated from AutoXAI4Omics for example task: *Multi-class classification in human transcriptomics.*** This is the default table generated as output in folder “Results” file name: scores__performance_results_testset.csv. Train refers to the training dataset for each ML model and Test refers to the held-out test dataset used for model evaluation.

| **model** | **accuracy_score_Train** | **accuracy_score_Test** | **f1_score_Train** | **f1_score_Test** | **f1_score_PerClass_Train** | **f1_score_PerClass_Test** |
| --- | --- | --- | --- | --- | --- | --- |
| **RandomForestClassifier** | 0.9924812030075190 | 0.9705882352941180 | 0.9924826777151760 | 0.9701525054466230 | [0.99029126 1.         0.98989899] | [1.         0.93333333 0.96296296] |
| **AdaBoostClassifier** | 1.0 | 0.9411764705882350 | 1.0 | 0.9407407407407410 | [1. 1. 1.] | [0.92307692 0.93333333 0.96296296] |
| **KNeighborsClassifier** | 0.9774436090225560 | 0.9411764705882350 | 0.9774436090225560 | 0.9417086834733890 | [0.97029703 1.         0.97029703] | [0.96       0.93333333 0.92857143] |
| **AutoXGBoost** | 1.0 | 0.8823529411764710 | 1.0 | 0.8769182886829950 | [1. 1. 1.] | [0.85714286 0.76923077 0.96296296] |
| **AutoLGBM** | 1.0 | 0.9411764705882350 | 1.0 | 0.9417086834733890 | [1. 1. 1.] | [0.96       0.93333333 0.92857143] |
| **AutoSKLearn** | 0.9924812030075190 | 0.7941176470588240 | 0.9924826777151760 | 0.785171568627451 | [0.99029126 1.         0.98989899] | [0.66666667 0.93333333 0.8125    ] |
| **AutoKeras** | 0.9774436090225560 | 0.8823529411764710 | 0.9774436090225560 | 0.883461210571185 | [0.97029703 1.         0.97029703] | [0.86956522 0.93333333 0.86666667] |
